# Long-term seasonality of marine photoheterotrophic bacteria reveals low cohesiveness within the different phylogroups

**DOI:** 10.1101/316059

**Authors:** Adrià Auladell, Pablo Sánchez, Olga Sánchez, Josep M. Gasol, Isabel Ferrera

**Affiliations:** Department of Marine Biology and Oceanography, Institut de Ciències del Mar (ICM-CSIC), Barcelona, Catalunya, Spain; Department of Genetics and Microbiology, Universitat Autònoma de Barcelona (UAB), Bellaterra, Catalunya, Spain; Centre for Marine Ecosystems Research, School of Science, Edith Cowan University, Joondalup, WA, Australia

## Abstract

Aerobic anoxygenic phototrophic (AAP) bacteria play a relevant role in the marine microbial food web, but little is known about their long-term seasonal dynamics. Using Illumina amplicon sequencing of the *puf*M gene coupled with multivariate, time series and co-occurrence analyses we examined their temporal dynamics over a decade at the Blanes Bay Microbial Observatory (NW Mediterranean). Phylogroup K (*Gammaproteobacteria*) was the most abundant over all seasons, with phylogroups E and G (*Alphaproteobacteria*) being often abundant in spring. A clear seasonal trend was observed in diversity, with maximum values in winter. Multivariate analyses showed sample clustering by season, with a relevant proportion of the variance (38%) explained by day length, temperature, salinity, phototrophic nanoflagellate abundance and phosphate concentration. Time series analysis showed that only 42% of the Amplicon Sequence Variants (ASVs) analyzed presented marked seasonality but these represented most of the abundance (92%). Interestingly, distinct temporal dynamics were observed within the same phylogroup and even within different ASVs conforming the same Operational Taxonomic Unit (OTU). Likewise, co-occurrence analysis highlighted negative associations between various ASVs within the same phylogroup. Altogether our results picture the AAP assemblage as highly seasonal, containing ecotypes with distinctive niche partitioning rather than being a cohesive functional group.

## Introduction

The development of high-throughput sequencing (HTS) has allowed the study of marine microbial diversity at an unprecedented scale (Sogin *et al.*, 2006). Mapping microbial communities in hundreds of samples from recent global expeditions has resulted in a more comprehensive picture of how they vary across space, which are the dominant bacterial groups and rare species, as well as which are the main controlling environmental parameters structuring these communities (Yooseph *et al.*, 2007; Salazar *et al.*, 2015; Sunagawa *et al.*, 2015). Seemingly important is determining the temporal dynamics of marine microbiomes. Disentangling community seasonality is crucial to unveil patterns of biodiversity, stability, predictability, resource preferences, interactions between species and response to disturbances, including global change. Long-term microbial observatories are thus key to understand microbial variation over time and across environmental gradients, particularly in temperate zones encompassing meteorological seasons (Kane, 2004; Buttigieg *et al.*, 2018).

To date, different seasonal studies conducted in fixed stations in the Atlantic and Pacific Oceans (i.e., the Western English Channel time series or the San Pedro Ocean Time Series (SPOT)) seem to concur that plankton turnover is related to meteorological seasons suggesting that community changes are driven by the environment, and that these patterns are repeatable over time (Gilbert *et al.*, 2012; Fuhrman *et al.*, 2015). Most of these studies have focused on determining the variation of phylogenetically distinct groups based on 16S or 18S rRNA gene sequencing for bacterioplankton and eukaryotic plankton respectively (Kim *et al.*, 2014; Fuhrman *et al.*, 2015; Martin-Platero *et al.*, 2018). However, we know that phylogenetic units based on ribosomal sequences may include different ecotypes given that closely related or even identical rRNA gene-identified species can possess different functional traits (Martiny *et al.*, 2013). Processes such as horizontal gene transfer (HGT) can disconnect functional from phylogenetic diversity (Louca *et al.*, 2016). A functional group particularly interesting is that of the polyphyletic (i.e., derived from more than one common ancestor through HGT) aerobic anoxygenic phototrophic (AAP) bacteria. For example, the *Roseobacter* clade, a major group of bacteria inhabiting surface ocean waters, contains both photoheterotrophic and strictly heterotrophic members (Koblížek *et al.*, 2013, 2015). While a considerable amount of information about the seasonality of microbial community structure and of some phylogroups (i.e.Galand *et al.*, 2010) exist, surprisingly little is known about the seasonality of specific functional groups, including the AAPs.

The AAPs have the ability of photoheterotrophy, that is, they are capable of using both organic matter and light as energy sources (Koblížek, 2015). Their discovery challenged previous simplistic views of the structure of ocean microbial food webs (Fenchel, 2001). AAP bacteria are relatively common in the euphotic zone of the oceans (Lami *et al.*, 2007; Yutin *et al.*, 2007; Cottrell and Kirchman, 2009), exhibit faster growth rates than other bacterioplankton groups (Ferrera *et al.*, 2011, 2017) and their cells are in general larger than most marine heterotrophic bacteria (Sieracki *et al.*, 2006). Altogether, these characteristics make them a relevant component of the microbial food web in processing organic matter, accounting for up to 25% of bacterial production (Hojerová *et al.*, 2011).

Phylogenetically, the AAPs are classified into 12 distinct phylogroups (from A to L) based on the structure of the *puf* operon and the *puf*M gene phylogeny (Yutin *et al.*, 2007), belonging to the *Alpha*-, *Beta*- and *Gammaproteobacteria* classes. Several studies have provided clues on their diversity and community structure in relation to environmental gradients across spatial scales using the *puf*M gene variability (Yutin *et al.*, 2007; Ritchie and Johnson, 2012; Lehours *et al.*, 2018). Contrarily, few studies have tackled their temporal dynamics. Ferrera *et al.* (2014) conducted a pioneer study in the NW Mediterranean using *puf*M HTS. The results indicated that the AAP assemblages are highly dynamic and undergo seasonal variations. Likewise, later results from the East coast of Australia concurred on identifying an important community structuring role of seasonality (Bibiloni-Isaksson *et al.*, 2016). Nevertheless, these studies were conducted over short time periods (1-year) but long-term studies are needed to unveil whether the observed seasonal trends are repeatable over time, as it has been shown to be the case for phylogenetic units based on ribosomal genes. Moreover, these studies have used the OTU approach, but the appearance of *threshold free* algorithms for sequence variants detection currently allows the analysis of ecotypes at a more refined level (Eren *et al.*, 2015; Callahan *et al.*, 2016), allowing to surpass the similarity clustering of 94% typically used for *puf*M gene.

Here, we present the first long time series exploration of marine AAP assemblages using Illumina sequencing and ASV analysis of the amplified *puf*M gene from monthly samples over 10 years at the coastal Blanes Bay Microbial Observatory (BBMO) in the NW Mediterranean Sea. The temporal patterns and predictability of the assemblage, as well as the long-term interactions between the different phylogroups have been explored, and the main environmental drivers acting upon the observed patterns identified. These analyses ultimately allow us to explore the level of ecological consistency within the different phylogenetic clades, that is, whether the different phylotypes of AAPs are ecologically cohesive or, contrarily, each phylogroup includes organisms showing niche partitioning.

## Material and Methods

### Location and sample collection

Surface waters were collected monthly at the Blanes Bay Microbial Observatory (41°40’N, 2°48’E), a shallow (∼20 m) coastal site about 1 km off the NW Mediterranean coast, as described elsewhere (Ferrera *et al.*, 2014). A total of 120 samples, from January 2004 to December 2013 were collected, and several environmental parameters were measured alongside sample collection (< 200μm). The sample code is as follows: BL + year (2 digits) + month + day (e.g BL110607). A matrix of environmental drivers was constructed using data of the following variables: day length, temperature and salinity measured with a CTD probe (model SAIV A/S SD204), Secchi depth, the concentration of inorganic nutrients determined spectrophotometrically using an Alliance Evolution II autoanalyzer according to standard procedures (Grasshoff *et al.*, 1983), chlorophyll *a* (Chl *a*) concentration (<200 μm fraction) measured from acetone extracts by fluorometry, and the abundances of heterotrophic prokaryotes, phototrophic prokaryotes (*Prochlorococcus* and *Synechococcus)* and eukaryotes measured by flow cytometry as described in Gasol and Morán (2015). Additionally, the abundance of *Cryptomonas*, *Micromonas*, phototrophic and heterotrophic nanoflagellates (PNF and HNF) was enumerated by epifluorescence microscopy from 4,6- diamidino-2-phenylindole (DAPI) stained samples, and bacterial activity was estimated from the incorporation of tritiated leucine (Smith and Azam, 1992). A total of 21 biotic and abiotic variables were used for statistical analysis, including those abovementioned as well as three different subgroups of PNF (the total, and fractions 2-5 μm and >5 μm), two of heterotrophic prokaryotes (high nucleic acid content, HNA, and low nucleic-acid content, LNA), and flow cytometrically determined picoeukaryotes group I (Peuk1; a discrete population distinguished in the cytogram characterized by relatively low scatter and fluorescence). Methods describing these variables are presented in Alonso-Sáez *et al*. (2008) and Gasol *et al*. (2016). The astronomical seasons (based on equinoxes and solstices) were used for establishing spring, summer, autumn and winter periods.

### DNA extraction, PCR amplification, sequencing and sequence processing

About 10 L of 200 μm pre-filtered surface seawater were sequentially filtered through a 20 μm mesh, a 3 μm pore-size polycarbonate filter (Poretics) and a 0.2 μm Sterivex Millipore filter using a peristaltic pump. Sterivex units were filled with 1.8 mL of lysis buffer (50 mM Tris-HCl pH 8.3, 40 mM EDTA pH 8.0 and 0.75 M sucrose) and kept at −80°C until extraction was performed using the phenol-chloroform protocol described by Massana *et al.* (1997). Partial amplification of the *puf*M gene (∼245 bp fragments) was done as described in Ferrera *et al.* (2014)using primers *puf*MF forward (5’-TACGGSAACCTGTWCTAC-3’, Béjà *et al.*, 2002) and *puf*WAW reverse (5’- AYNGCRAACCACCANGCCCA-3’,Yutin *et al.*, 2005). Sequencing was performed in an Illumina MiSeq sequencer (2 *x* 250 bp, Research and Testing Laboratory; http://rtlgenomics.com/). Primers and spurious sequences were trimmed using *cutadapt* (Martin, 2011) trimming ~50 bp. DADA2 was used to differentiate exact sequence variants (Callahan *et al.*, 2016). DADA2 resolves ASVs by modeling the errors in Illumina-sequenced amplicon reads. The approach is *threshold free*, inferring exact variants up to 1 nucleotide of difference using the Q scores in a probability model. For comparison purposes, samples were processed with an UPARSE pipeline (Logares, 2017) generating OTUs clustered at 94% of sequence similarity, typically used for the *puf*M gene (Ferrera *et al.*, 2014; Bibiloni-Isaksson *et al.*, 2016). The OTU/ASV correspondence was calculated with the number of nucleotide differences between them, considering <12 the same OTU. Each ASV sequence was classified taxonomically using a custom-made database of the *puf*M gene created combining previous studies (Yutin *et al.*, 2007; Lehours and Jeanthon, 2015) and other *puf*M sequences from the GenBank database. Assignations with identity <80% and alignment <170 bp were considered unclassified. Sample BL120313 (March 2012) was discarded due to low quality criteria.

### Quantitative Polymerase Chain Reaction

The relative abundance of AAP was estimated by quantitative polymerase chain reaction (qPCR) of the marker gene *puf*M, as described in Ferrera *et al.*, (2017b). Reactions were performed in triplicate on a MyiQ™ Single-Color Real-Time PCR Detection System (Bio-Rad) using Maxima SYBR Green qPCR Master Mix (2X; Fermentas). Standard curves were generated from amplification of *Roseobacter* sp. COL2P genomic DNA (Koblížek *et al.*, 2010). We remark that we did not conduct absolute but rather relative quantifications since our goal was only to test whether a seasonal trend existed.

### Statistical analyses

All analyses were performed using the R language, with *phyloseq* and *vegan* packages (McMurdie and Holmes, 2013; Oksanen *et al.*, 2013; R Core Team, 2014). Alphadiversity was analyzed using the Chao1, Shannon and Simpson indices. Betadiversity was analyzed using a Bray-Curtis dissimilarity matrix from log10 + 1 transformed data. We used distance-based Redundancy Analysis (dbRDA,Legendre and Legendre, 1988) to find the environmental predictors (with a previous scaling) that best explained the patterns of abundance and diversity of AAPs over time, with a previous multivariate non-parametric ANOVA for selecting significant variables (*p*<0.01). A time-decay analysis of the assemblage was computed excluding rare ASVs as recommended elsewhere (Faust *et al.*, 2015). ASVs were considered rare when presenting always less than 1% of relative abundance, following the description by Alonso-Sáez *et al.* (2015a).

### Time series analysis

Fourier time series analysis was performed to study the AAP assemblage dynamics over a decade. An interpolation of the discarded sample (BL120313) was used to maintain equidistant time points. Relative abundances were transformed logarithmically adding a pseudocount of 1. A Fisher G-test from the R package *GeneCycle* was used to determine the significance of the periodic components, with a threshold of *p*<0.01 (Ahdesmaki *et al.*, 2015). The time series was decomposed in three components - the seasonal periodicity (oscillation inside each period), the trend (evolution over time) and residuals-through local regression by the *stl* function. Additionally, the autocorrelogram was calculated through the *acf* function.

### Network construction

Many methods exist to identify associations between OTUs/ASVs and environmental variables (Weiss *et al.*, 2016). However, microbiome datasets present some difficulties due to the sparsity (presence of a large number of zeroes) and compositionality (relative abundances subedited to a simplex, like 100%) (Kurtz *et al.*, 2015; Gloor *et al.*, 2017). Since our study spans 10 years, codependency also exists, limiting the possible methods to use, and among those we chose Local Similarity Analysis (LSA) (Xia *et al.*, 2011; Durno *et al.*, 2013). Briefly, given a time series data and a delay limit, LSA finds the configuration of the data that yields the highest local similarity (LS) score. We applied the Aitchison log-centered ratio transformation, more adequate for compositional data (Gloor *et al.*, 2017). Only the ASVs present in >5 samples and the environmental variables presenting <5% of missing values were used. The remaining missing values were replaced by imputation with the *mice* package (Azur *et al.*, 2012). Only interactions with a LS >0.6, significance of p<0.001 and 1- month delay were taken into account. The network was plotted using the *ggraph* package (Pedersen, 2017), including the nodes - ASVs and environmental variables - and the edges (connections between nodes) derived from the algorithm scores.

### Reproducibility

The code and details used for preprocessing and statistical analyses can be found in the Gitlab repository: https://gitlab.com/aauladell/AAP_time_series. Sequence data has been deposited in Genbank under accession number PRJNA449272.

## Results and discussion

Interpretations of observations that use genes as biological objects depend to a large extent on similarity thresholds that may group together an appreciable amount of phenotypic diversity. With the appearance of *threshold free* algorithms for the detection of sequence variants, the analysis of ecotypes can be applied to a more refined level (Eren *et al.*, 2015; Callahan *et al.*, 2016). In that way, we were able to divide the *puf*M OTUs and distinguish ASVs showing a divergent seasonal behavior potentially representing distinct ecotypes. The decadal analysis allowed determining whether the community structure trends are robust, as well as to explore if different ASVs present similar seasonal behavior to, ultimately, test the ecological cohesiveness of these populations, a result only possible through long-term time series analysis.

### Community composition and structure

While the number of OTUs detected was of 449 (94% similarity cutoff), the number of ASVs was 820. Of these, 276 present only 1 nucleotide variation between sequences. Rarefaction curves showed asymptotes at the sequencing depth for most samples (Figure S1), suggesting that our dataset covers most of the AAP diversity in Blanes Bay. In comparison with the number of OTUs observed in previous temporal studies (82 OTUs, Ferrera *et al.*, 2014, and 89 Bibiloni-Isaksson *et al.*, 2016), our study presents a more complete picture of the *puf*M diversity, being the largest dataset of AAP diversity ever reported. Alphadiversity measurements showed that values were higher during winter (mean 51, max 126 observed ASVs), decreasing to minimum values in the spring-summer period, specifically during May-August (mean 35, max 77) (Figure 1). The differences between winter and spring/summer were statistically significant (ANOVA, *p*<0.05). A similar trend was observed when computing Shannon and Simpson diversity indices (Figure 1).

**Figure 1.**
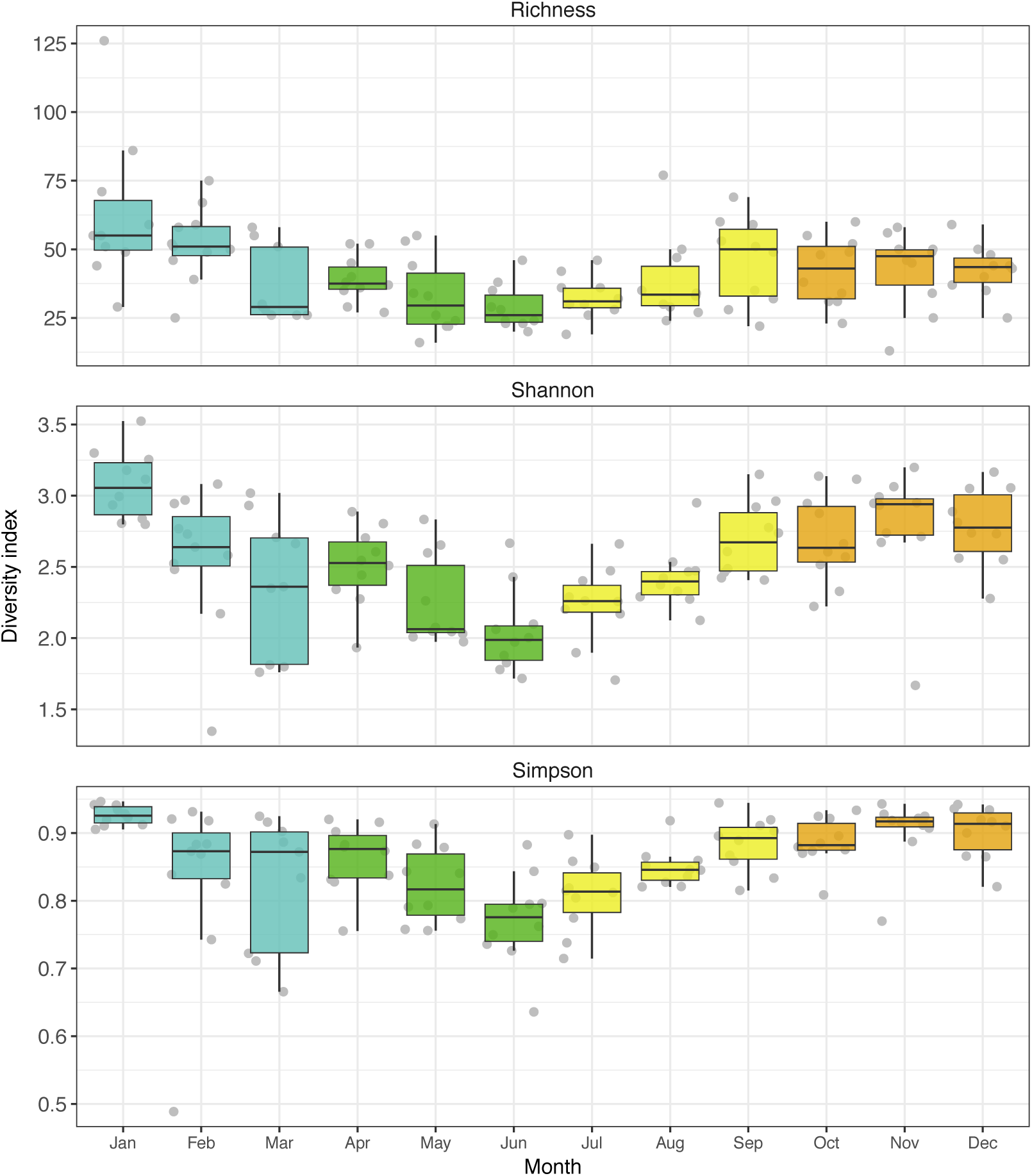
Alphadiversity distribution of the AAP community for each month over the studied decade (2004-2013). Number of observed ASVs, Shannon and Simpson indexes are shown colored by season in the top, middle and bottom panels respectively. Each boxplot presents the average and the 25% and 75% limits with the distribution of 10 data points in grey (with the exception of March, with 9 data points).

We observed a negative correlation between day length and the Shannon index (N=119, Pearson r=- 0.57, *p*<0.01). The diversity relationship with day length had been observed previously in long-term bulk bacterioplankton community studies (Gilbert *et al.*, 2012), as well as when targeting specific phylogenetic groups such as the SAR11 (Salter *et al.*, 2015). A possible explanation is that the deep winter mixing allows the development of high diversity assemblages in contrast to the selection of specific oligotrophic ecotypes occurring during the stratified summer season (Salter *et al.*, 2015). Our results confirm that the trend observed for 16S rRNA-defined taxa is comparable to that of the AAP functional assemblage. Interestingly, the trend in alphadiversity is contrary to that of AAP abundance (Figure S2). An abundance increase during summer as compared to winter and fall (*p*<0.01) was measured by qPCR. These results support the observations from other studies that also presented the AAPs to be more abundant during summer (Ferrera *et al.*, 2014; Bibiloni-Isaksson *et al.*, 2016).

Regarding community composition and structure across the decadal period (Figure 2, Figure S3), phylogroup K (*Gammaproteobacteria*) was the most ubiquitous and abundant over the years (83.4 ± SE 2.2, mean relative abundance). Contrarily to observations for other oceanographic regions (Yutin *et al.*, 2007), gammaproteobacterial AAPs appear to be the most abundant in the Mediterranean Sea (Lehours *et al.*, 2010; Ferrera *et al.*, 2014). Yet, February and March showed a decrease in their contribution (59.6% and 52% mean respectively). During these months, phylogroups E and G (*Rhodobacter* and *Roseobacter*-like, respectively) increased in their relative contribution albeit with a high variation over the decade (~ ±14% and ±25% SD respectively). As an example, the contribution of phylogroup E in 2006 reached ~47% of the total community sequences from February to April. Conversely, in the same period of 2005 the contribution of this group was really low (2%). Overall, phylogroups F (*Rhodobacterales*-like), H (uncultured), J (*Rhodospirillales-*like) and the unclassified ASVs presented a mean relative contribution below 1%. However, these groups presented occasional peaks (>1% relative abundance) with no clear periodic trend. For example, phylogroup J showed a contribution of 14% in February 2012 (Figure S3). When analyzing the structure of the assemblage, we observed that most reads corresponded to very few sequence variants (Figure S3). Thus, the structure of AAPs follows a comparable pattern to the whole microbial communities, i.e., they are dominated by a few species while containing a large number of species represented by only a few individuals (Pedrós-Alió, 2012).

**Figure 2.**
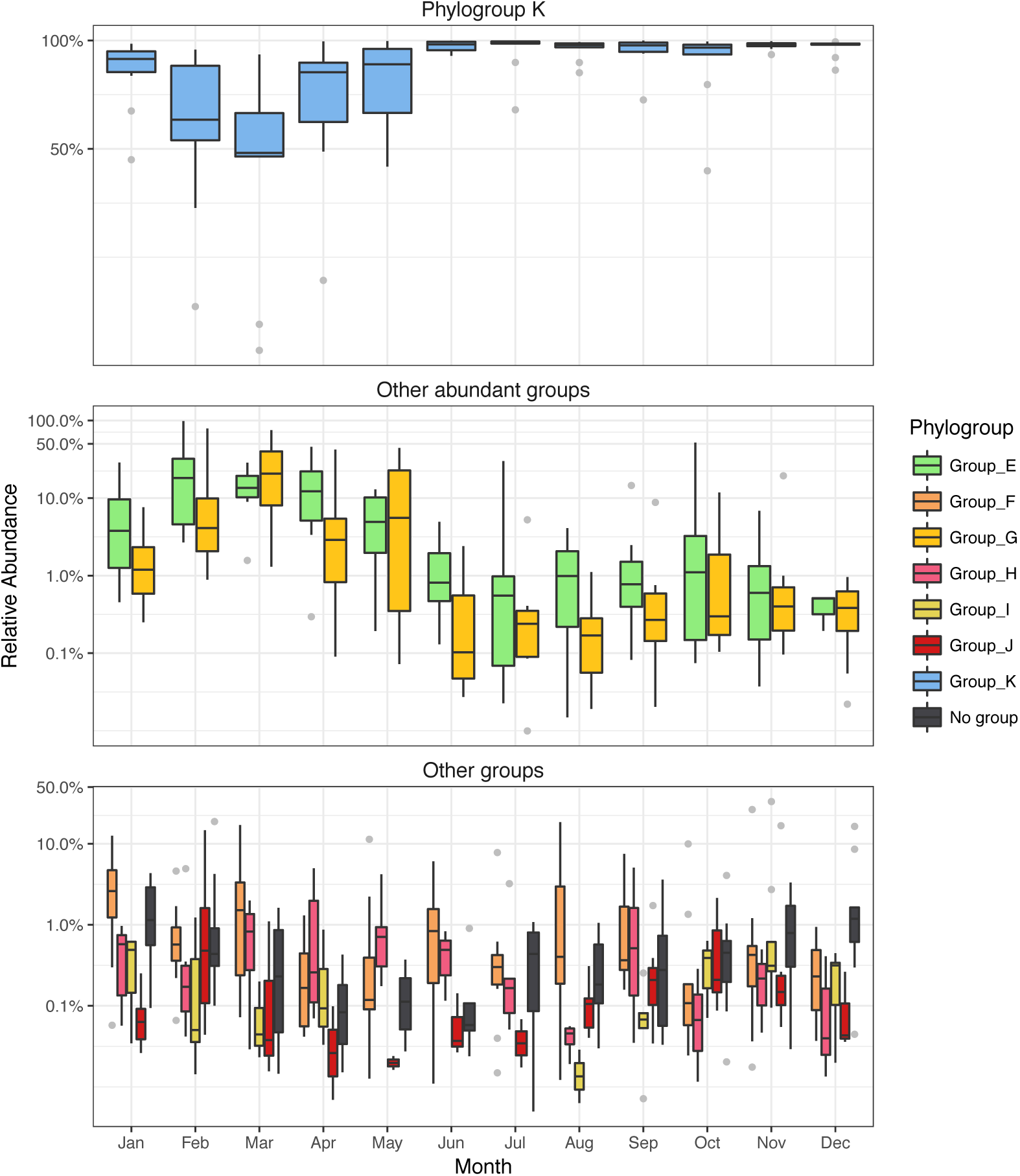
Variation in the relative abundances of phylogroups K (top panel), G, E (middle panel) and phylogroups F, H, J and the unclassified group (bottom panel) for each month over the studied decade (2004-2013). Each boxplot presents the average and the 25% and 75% limits with the distribution of 10 data points in grey (with the exception of March, with 9 data points).

Within the *Gammaproteobacteria*, the high abundance values of phylogroup K (represented by sequences affiliated to the type strain *Congregibacter litoralis* of the NOR5/OM60 clade) were related only to a few sequence variants (10-15 depending on the year). The most abundant sequence variant was ASV1, which corresponded to 18% of the total read abundance, followed by ASV2 (8.2%), ASV5 (6.6%) and ASV6 (6.2%). A similar trend was observed for phylogroups E (ASV8, 5.6%) and G (ASV14, 3.7%), both *Rhodobacterales*-like. The results were even more pronounced at the OTU level: a single OTU corresponded to 35% of the total read abundance. A previous study on 16S rRNA gene diversity in the same location suggested that *Alphaproteobacteria* dominates the bacterial assemblages during the spring bloom that occurs in that study site (Alonso-Sáez *et al.*, 2007). Likewise, in the 1-year study of AAP diversity conducted byFerrera *et al.* (2014), alphaproteobacterial members of the AAP were more abundant during this season. In our long-term analysis, despite we also observed that in some years phylogroups G, E and F were more abundant during March-May (Figure 2), taking into account the entire decade, phylogroup K was overall the most abundant during these months (65% mean relative abundance).

### Betadiversity analysis

To determine similarities between samples, betadiversity was compared using Bray-Curtis dissimilarity and non-metric multidimensional scaling (nMDS). The nMDS indicated a clear separation of the samples at different temporal scales: by month (PERMANOVA R^2^=0.39, *p*<0.001) and by season (PERMANOVA R^2^=0.25, *p*<0.001). This structure was maintained using alternative distance measurements (Figure S4). Spring and winter samples presented higher dissimilarity within their values in comparison with summer and autumn (Fig S4). The reasons for this pattern are uncertain but it could be related to higher environmental variability or to the mixing of the water column that occurs during winter (Gasol *et al.*, 2016). Noteworthy, despite the monthly assemblages seemed to be recurrent over time there was a remarkable exception. In November 2006, the ASV distribution was completely different to that found during that month in other years. In that particular sample, betaproteobacterial phylogroup I (33%), and alphaproteobacterial phylogroups F (26%) and G (19%) dominated the composition of the community (Figure S5). When observing all the environmental variables for that sample, we found that particularly the abundances of picoeukaryotes and phototrophic nanoflagellates were above seasonal averages (Figure S6). This observation is a clear example on how members typically found in the rare biosphere can eventually become abundant (Pedrós-Alió, 2012).

Community structure distribution was significantly linked to day length, temperature, salinity, PO_4_^3-^ concentration and the abundance of phototrophic nanoflagellates as revealed by distance-based redundancy analysis (Figure 3, PERMANOVA *p*<0.01). With these 5 variables, the dbRDA explained 38.1% of the variation, with the 2 first axis explaining the 32.9%. Interestingly, a group of samples, mainly from winter and spring, appeared to be related to the phototrophic nanoflagellate abundances (Figure 3), and to ASVs belonging to phylogroups E and G (*Rhodobacterales*-like) (Figure S7). This biotic parameter could be related to the phytoplankton spring bloom that typically occurs in February-March in Blanes (Gasol *et al.*, 2016). Late spring and early summer samples were mostly influenced by day length and temperature, with autumn samples partially influenced by salinity. Day length has previously been shown to explain the seasonal variability of the whole bacterioplankton (Gilbert *et al.*, 2012) and AAP community structure (Ferrera *et al.*, 2014), but the mechanisms underlying this relationship are unclear. Summing up, the dissimilarity measurements with the environmental variables as explanatory variables distinguished 3 groups: the summer samples, related to the high abundance of gammaproteobacterial ASV1 and temperature, the fall/early-winter cluster, with more diverse communities related to other gammaproteobacterial ASVs and salinity, and the winter-spring samples, highly variable due to the effect of diverse ASVs related to alphaproteobacterial phylogroups G and E and to the phototrophic nanoflagellate abundance.

**Figure 3.**
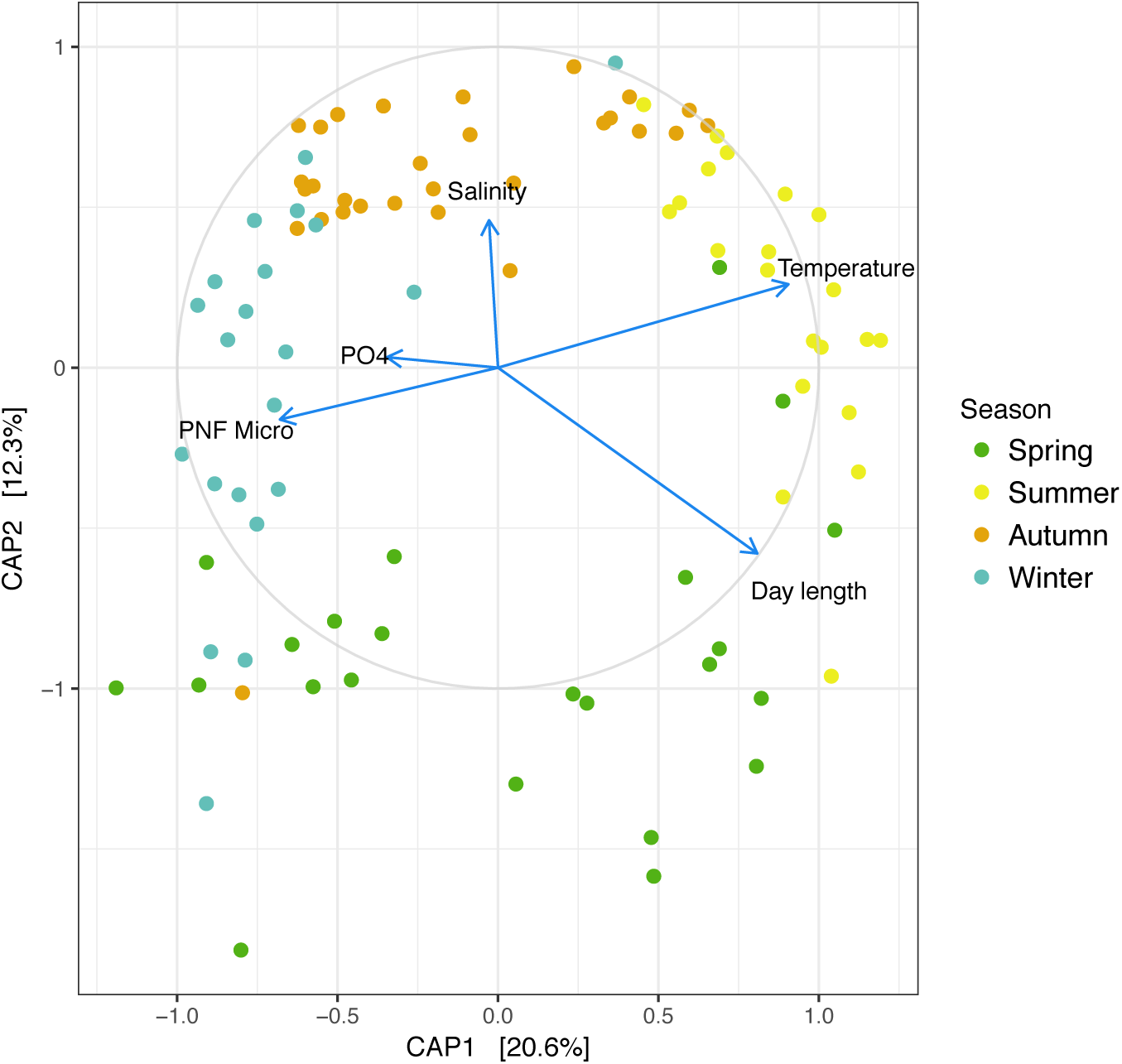
Distance based redundancy analysis of the samples (dots) with the 5 explanatory variables influencing the distribution (PERMANOVA *p*<0.01; day length, temperature, salinity, phosphate concentration (PO_4_^-3^) and phototrophic nanoflagellate abundance (PNF)) shown in arrows. The ordination was performed on the Bray-Curtis dissimilarity of log_10_ transformed data (with a pseudocount of 1) matrix. Each sample is colored by season.

Finally, to study the evolution of the community over the decade, a comparison of the Bray-Curtis similarity between samples was plotted against the time lag, commonly known as the time-decay curve (Shade *et al.*, 2013; Fuhrman *et al.*, 2015). In our study, the assemblage was maintained with a median similarity of 0.45, with oscillations with the maxima every 12 months (~0.55) and the minimum every 6 months (~0.39). The similarity oscillates with one-year apart peaks, indicating a high seasonal behavior, with a slight negative slope in the global linear regression (Figure 4). To our knowledge, this approach had not yet been applied to a functional group defined by a marker gene. Comparing the results to the 16S rRNA data from the SPOT and the Western Channel time series results (see Hatosy *et al.*, 2013 for details) we observe that in SPOT, the seasonal turnover is less clear than in our location, and an initial decay of similarity is observed, reaching a plateau of similarity after 4 years (Fuhrman *et al.*, 2015). In the Western Channel, the seasonality is equally marked but the decay is more pronounced. A possible explanation for the differences is that our comparison accounts only for a highly seasonal sub-community while the overall bacterial/prokaryotic community varies more. Further analyses with other functional genes would help understand how the different functional groups distribution varies over time.

**Figure 4.**
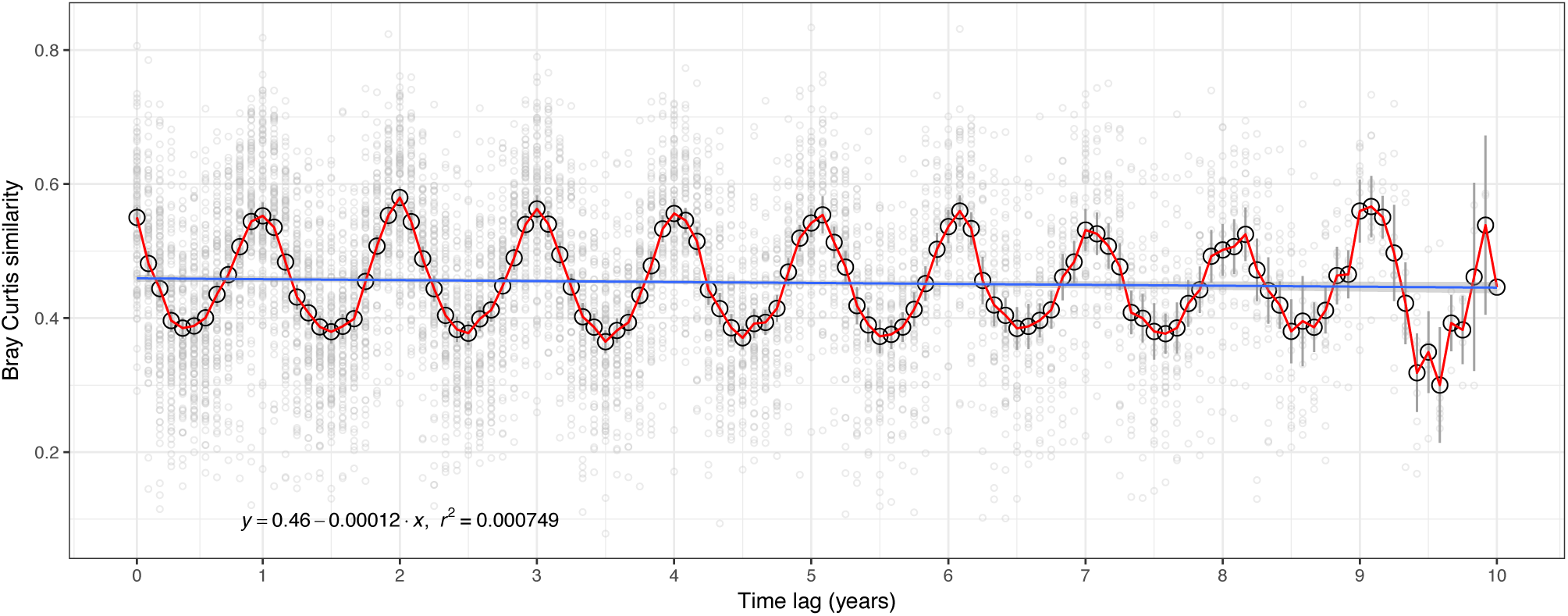
Bray-Curtis similarity between samples plotted against the time lag between each of them (time-decay plot). Mean similarity values for each lag are plotted in an empty black dot with confidence interval bars (background grey filled dots show each comparison). A linear regression is plotted in blue, with the standard error around the prediction.

### Time series analysis

Through long-term analysis we approached the seasonality of each ASV by evaluating if their relative abundance distribution presented a significant periodicity (Fisher G-test), and if so, compared them at different levels: across closely related sequences (ASVs) and across sequence clusters (OTUs and phylogroups). Seasonal patterns were present in 58 (*p*<0.01) out of 127 ASVs analyzed (those present in >5 samples), predominantly affiliated with phylogroups K (39 ASVs), E (5) and G (3) and the unclassified group (7) (Table 1, Table S1). These seasonal ASVs corresponded to 92% of the total read counts, and 81% of the counts corresponded to phylogroup K (*Gammaproteobacteria*). All periodicities found were of 1 year, with the exception of ASV152 (*Gammaproteobacteria*), presenting a periodicity of 2 years. Some of these ASVs were always abundant (>1%) regardless of season (all from phylogroup K), some presented values above 1% in a specific season (seasonally abundant), and other ASVs peaked in abundance (>1%) only occasionally (i.e. opportunistic; see examples in Figure S8, S9, S10, S11). In fact, most ASVs were opportunistic, with low abundance during the decade and peaking only occasionally. Various studies of the whole bacterioplankton community have presented these variety of strategies coexisting within a given clade (Shade *et al.*, 2014; Alonso-Sáez *et al.*, 2015; Fuhrman *et al.*, 2015). Our results reveal that this trend is maintained for a specific functional assemblage, with few generalist ecotypes and a larger pool of specialized ASVs within each phylum. While the previous AAP temporal studies provided insights of the inter-annual community structure, only long-term analyses can differentiate tendencies from occasional peaks of rare specialized ecotypes.

**Table 1.**
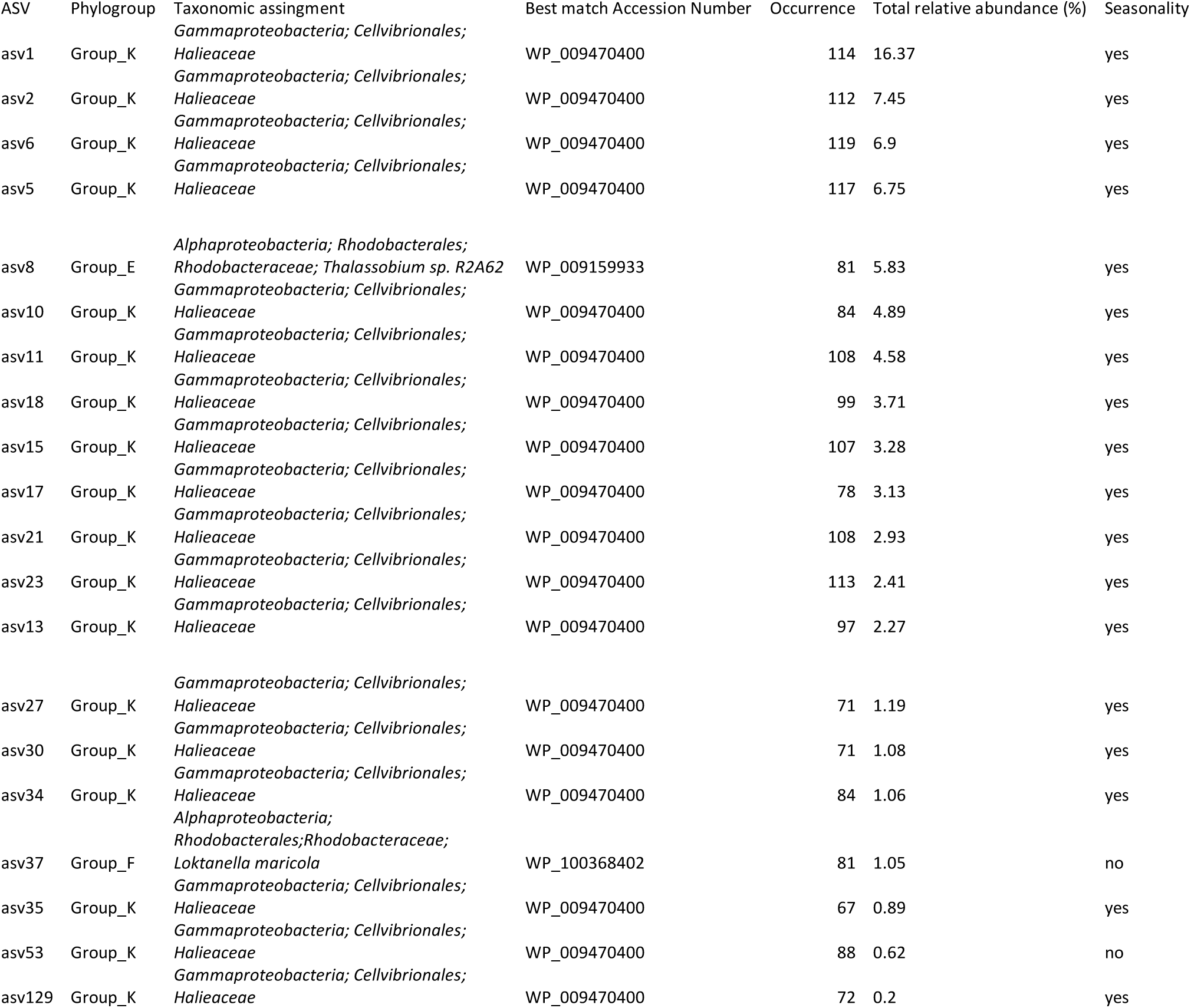
Summary information for the top 20 ASVs (based on abundance). The following columns are listed: ASV name, phylogroup correspondence, taxonomy based on Blastx against the NCBI RefSeq (release 87) database, occurrence (number of samples in which they were present being the maximum number 119), mean relative abundance, seasonality behavior, and month of maximum mean relative abundance.

Comparing among the seasonal ASVs, we distinguished different behaviors. For example, ASVs divergent enough to form distinct OTUs but belonging to the same phylogroup (Figure 5A) did not always follow the same distribution; e.g. for phylogroup K, the annual maxima of ASV1 occurred during June and July with a minimum in February/March, whereas ASV10 presents the opposite distribution. Other phylogroup K ASVs, like ASV5, presented a less marked seasonality, with a maximum in September. Contrarily, most ASVs belonging to phylogroups G and E followed a similar trend among them, with their maxima in March, being an exception ASV86, presenting a maximum in September.

**Figure 5.**
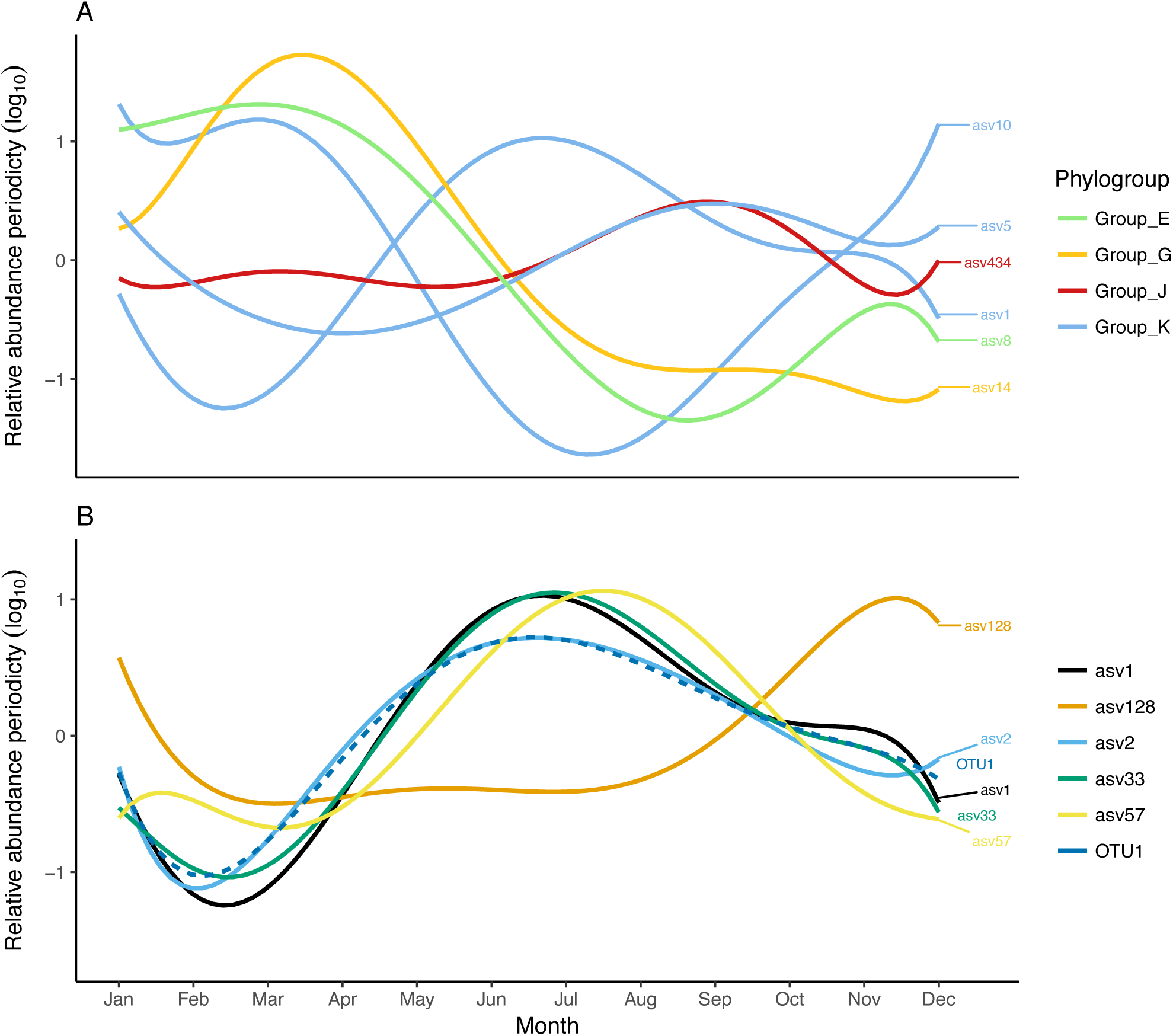
Seasonal component of the relative abundance distribution (log10 +1 transformed) for some remarked ASVs fitted with a polynomial function. (A) Various ASVs with distant nucleotide similarity, colored by phylogroup assignation. (B) Various ASVs belonging to the same OTU (nucleotide differences <12). The patterns were defined based on the relative abundance dynamics of 10 years (2004-2013) by time series analysis.

Further, comparing the seasonality of closely related ASVs (that would form the same OTU) we observed that, in general, these displayed similar temporal patterns but some notable exceptions exist. An example is represented in Figure 5B in which the seasonal periodicities of 5 closely related ASVs – all corresponding to OTU1, clustering at 94% - are plotted together. In this figure, a slight succession of the summer month maxima can be observed (ASV2 peaking slightly before ASVs 33 and 1, with ASV57 afterwards), being all these only 1 nucleotide different among them. Yet, ASV128 (distance of 4 nucleotides) presents a completely different distribution, with peaks during winter. The existence of divergent distributions of ASVs composing the same OTU demonstrates the need to break apart the clusters of related sequences, since these can hide interesting ecological patterns.

When establishing seasonality at the phylogroup level, we found that group K (*Gammaproteobacteria*) and J (*Rhodospirillales*-like, *Alphaproteobacteria*) did not present significant seasonal patterns (Figure S12, S13). Phylogroup J only presented 1 seasonal ASV but in the case of phylogroup K, the disparity of distributions of the various sequences results in the loss of a specific signal differentiable when computing seasonality at the group level. Contrarily, the autocorrelograms showed phylogroup G presenting a high value (max.0.38 over a year), followed by phylogroups E and I (Figure S12B). These results could indicate a higher degree of ecotype differentiation in gammaproteobacterial phylogroup K as compared to alphaproteobacterial phylogroups E and G in the NW Mediterranean.

Ferrera *et al.* (2014) suggested that the high abundance of *Gammaproteobacteria-*like AAPs in summer together with the relationship with day length could be an indication of phototrophy serving as auxiliary energy source to use the refractory dissolved organic carbon that accumulates in summer in this coastal station (Vila-Reixach *et al.*, 2012; Romera-Castillo *et al.*, 2013). Our results point out that this hypothesis may not be true for the whole community and rather only for certain ecotypes, such as ASV1, 2 and 5. Contrarily, ASV5, 11 and 21 possibly have preference for a more eutrophic environment like that of spring blooms, or are associated to specific phytoplankton species, presenting a high fluctuation likely because of the relationship with these events.

Lehours *et al*. (2018) recently tested the ecological consistency of the AAP across different oceanic regions and, interestingly, identified clades with good ecological and phylogenetic coherence. Our temporal analyses complement their spatial study and adds a new level of complexity by showing that, in some cases, even the ASVs from the same phylogroup and even those contained within the same OTU can present different distributions that could translate into different ecology. These analyses could be expanded adding genomic context with the assignation of sequence variants with Metagenome Assembled Genomes (MAGs) containing *puf*M genes with an exact match. Additionally, experimental work with cultures of these ASVs could shed light on the reasons for their high prevalence in the environment.

### Co-occurrence analysis of the assemblage

Out of the 127 ASVs present in >5 samples and 14 environmental variables, the final network presented 49 nodes and 54 edges, separated in 12 components (that is, 12 subnetworks without connections among them) with 6 of them presenting more than 3 nodes (Figure 6). Of these subclusters, some were composed by closely related ASVs (that would form the same OTU) and presented the same ecological distribution, like ASV1, 2 and 15 (all OTU1, group K). Others, like ASV33 and 63 (also OTU1), were dissociated, with differences in their distribution. The network revealed that temperature plays an important role on the distribution of some ASVs, with a lagged negative correlation with ASV11, 18 and 10 and a positive one with ASV27. We also observed that phylogroups G and E (both *Alphaproteobacteria*) are positively related between them while presenting negative associations with ASV30 and 35 (phylogroup K). The negative association of phylogroup K and phylogroup G OTUs highlighted in this study was previously spotted by Ferrera *et al.* (2014) as well as by Bibiloni-Isaksson *et al.* (2016). Genome sequencing and strain characterization of AAP isolates have revealed differences in the metabolic capabilities and physiological properties of different strains (Koblížek *et al.*, 2003; Fuchs *et al.*, 2007). The lack of seasonality at a higher taxonomic level for some phylogroups, as well as the network negative associations, may suggest that intergroup competition between different AAPs exists or that different AAPs possess diverse strategies to adapt to a changing environment.

**Figure 6.**
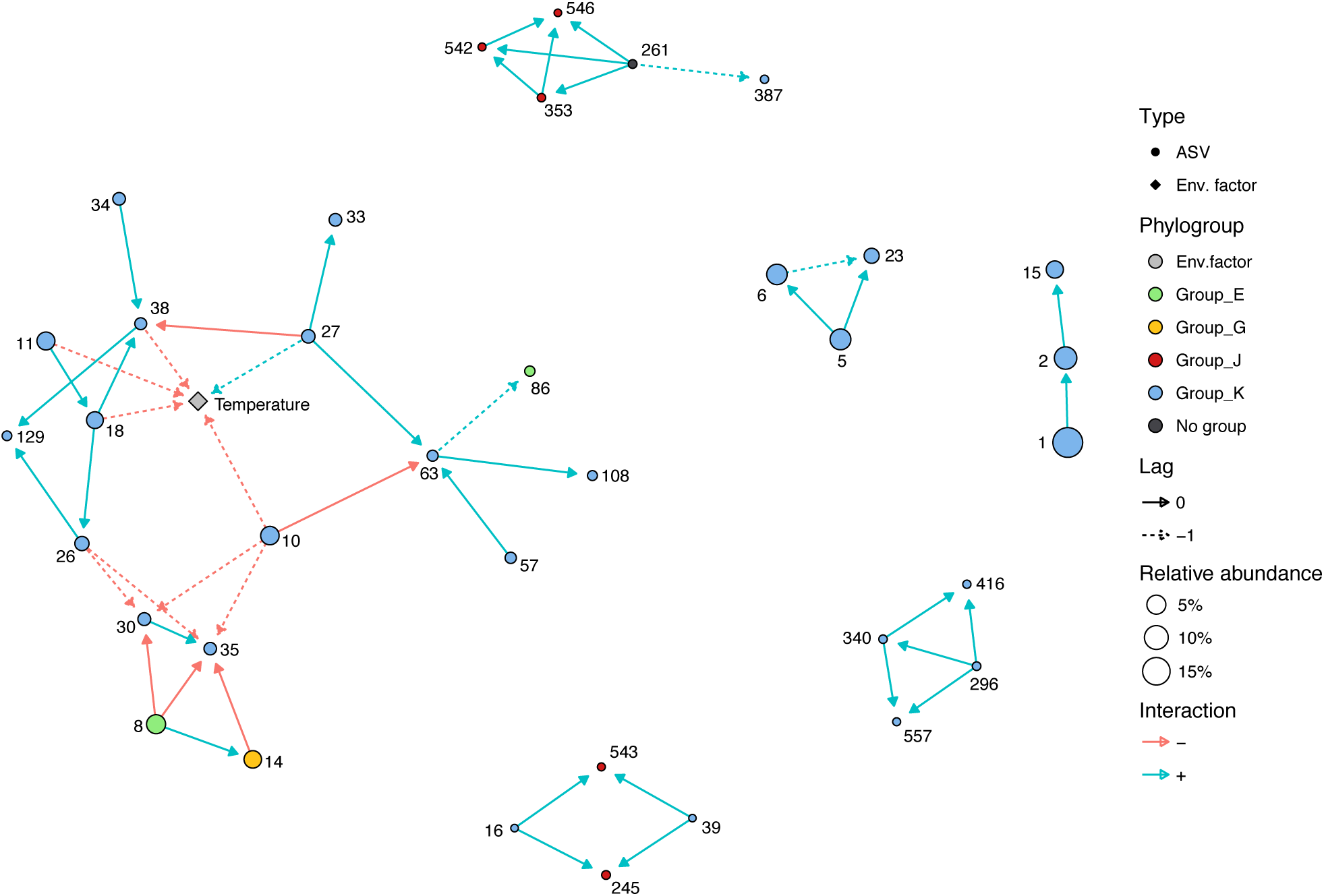
Network created with fast local similarity analysis. The network contains 37 nodes and 48 edges. Node shape is related to the type of variable, with the color specifying the phylogroup correspondence and size the total relative abundance. Edges can be lagged (discontinuous line) or direct and have negative (i.e., anticorrelation) or positive local scores (LS). The label on the nodes indicates the ASV number.

### Concluding remarks

Our study shows the advantages of long-term time series for describing the microbial seasonal periodicities of a specific functional group. The results confirm that the AAPs present the peak of diversity during winter, opposite to their abundance, and that gammaproteobacterial AAPs are the most abundant in the Mediterranean Sea year-round. Moreover, the study shows that the assemblages have recurrent seasonality over the years. Yet, despite the most abundant ASVs were clearly seasonal, phylogroup K as a group did not present a stable seasonality. Contrarily to the recent spatial study of Lehours *et al.* (2018)in which they reported ecological cohesiveness when comparing contrasting biomes, we found that the AAP do not appear as coherent when studying their temporal dynamics at a high-resolution level and seem to be instead formed by different ecotypes with distinctive niche partitioning. Overall, these results show that the analysis of a long time series allows exploring in-depth patterns of a highly dynamic microbial group and provides a framework for modeling their ecological role in relation to seasonality of the marine carbon cycling.

## Acknowledgements

We thank the many people involved in maintaining the BBMO and those taking care of sampling, particularly Clara Cardelús and Vanessa Balagué. We would also like to thank Ramon Massana and Irene Forn for providing microscopy counts. We thank the Marbits bioinformatics platform at ICM-CSIC, particularly Ramiro Logares, and Anders Kristian Krabberød (University of Oslo) for help with computing analyses. qPCR analysis was done at the Institute of Evolutionary Biology (Barcelona) thanks to José Luís Maestro. This research was funded by grants MALASPINA2010 (CSD2008-00077) from the former Spanish Ministry of Science and Innovation and REMEI (CTM2015-70340-R) from the Spanish Ministry of Economy, Industry and Competitiveness.

## Statement of authorship

I.F. and J.M.G. conceived the study; I.F. and O.S. designed and performed laboratory analyses; A.A. and I.F. analyzed the data; A.A. and I.F. wrote the paper and all authors commented and revised it.

---

**Supplementary Table 1.** Summary information of the 820 amplicon sequence variants. The following columns are listed: ASV, sequence, phylogroup, OTU correspondence, nucleotide differences, occurrence (number of samples present), total relative abundance (%), seasonality, month of maximum relative abundance, median (% for the specific maximum of relative abundance) and standard error (SE).

## References

Ahdesmaki M, Fokianos K, Strimmer K. (2015). Package ‘ GeneCycle ’. http://www.cs.tut.fi/~ahdesmak/robustperiodic/index.html.

Alonso-Sáez L, Balagué V, Sà EL, Sánchez O, González JM, Pinhassi J, et al. (2007). Seasonality in bacterial diversity in north-west Mediterranean coastal waters: Assessment through clone libraries, fingerprinting and FISH. FEMS Microbiol Ecol 60: 98–112.

Alonso-Sáez L, Díaz-Pérez L, Morán XAG. (2015). The hidden seasonality of the rare biosphere in coastal marine bacterioplankton. Environ Microbiol 17: 3766–3780.

Azur MJ, Stuart EA, Frangakis C, Leaf PJ. (2012). Multiple Imputation by Chained Equations: What is it and how does it work? Int J Methods Psychiatr Res 20: 40–49.

Béjà O, Suzuki MT, Heidelberg JF, Nelson WC, Preston CM, Hamada T, et al. (2002). Unsuspected diversity among marine aerobic anoxygenic phototrophs. Nature 415: 630–633.

Bibiloni-Isaksson J, Seymour JR, Ingleton T, van de Kamp J, Bodrossy L, Brown M V. (2016). Spatial and temporal variability of aerobic anoxygenic photoheterotrophic bacteria along the east coast of Australia. Environ Microbiol 18: 4485–4500.

Buttigieg PL, Fadeev E, Bienhold C, Hehemann L, Offre P, Boetius A. (2018). Marine microbes in 4D — using time series observation to assess the dynamics of the ocean microbiome and its links to ocean health. Curr Opin Microbiol 43: 169–185.

Callahan BJ, McMurdie PJ, Rosen MJ, Han AW, Johnson AJA, Holmes SP. (2016). DADA2: High-resolution sample inference from Illumina amplicon data. Nat Methods 13: 581.

Cottrell MT, Kirchman DL. (2009). Photoheterotrophic microbes in the arctic ocean in summer and winter. Appl Environ Microbiol 75: 4958–4966.

Durno WE, Hanson NW, Konwar KM, Hallam SJ. (2013). Expanding the boundaries of local similarity analysis. BMC Genomics 14: 1–14.

Eren AM, Morrison HG, Lescault PJ, Reveillaud J, Vineis JH, Sogin ML. (2015). Minimum entropy decomposition: Unsupervised oligotyping for sensitive partitioning of high-throughput marker gene sequences. ISME J 9: 968–979.

Faust K, Lahti L, Gonze D, de Vos WM, Raes J. (2015). Metagenomics meets time series analysis: Unraveling microbial community dynamics. Curr Opin Microbiol 25: 56–66.

Fenchel T. (2001). Marine bugs and carbon flow. Science (80- ) 292: 2444 LP-2445.

Ferrera I, Borrego CM, Salazar G, Gasol JM. (2014). Marked seasonality of aerobic anoxygenic phototrophic bacteria in the coastal NW Mediterranean Sea as revealed by cell abundance, pigment concentration and pyrosequencing of *puf*M gene. Environ Microbiol 16: 2953–2965.

Ferrera I, Gasol JM, Sebastián M, Hojerová E, Kobížek M. (2011). Comparison of growth rates of aerobic anoxygenic phototrophic bacteria and other bacterioplankton groups in coastal mediterranean waters. Appl Environ Microbiol 77: 7451–7458.

Ferrera I, Sanchez O, Kolarova E, Koblížek M, Gasol JM. (2017a). Light enhances the growth rates of natural populations of aerobic anoxygenic phototrophic bacteria. ISME J 11: 2391–2393.

Ferrera I, Sarmento H, Priscu J, Chiuchiolo A, Gonzalez JM, Grossart H-P. (2017b). Diversity and distribution of freshwater aerobic anoxygenic phototrophic bacteria across a wide latitudinal gradient. Front Microbiol 8: 175.

Fuhrman JA, Cram JA, Needham DM. (2015). Marine microbial community dynamics and their ecological interpretation. Nat Rev Microbiol 13: 133–146.

Galand PE, Gutiérrez-Provecho C, Massana R, Gasol JM, Casamayor EO. (2010). Inter-annual recurrence of archaeal assemblages in the coastal NW Mediterranean Sea (Blanes Bay Microbial Observatory). Limnol Oceanogr 55: 2117–2125.

Gasol JM, Cardelús C, Morán XAG, Balagué V, Forn I, Marrasé C, et al. (2016). Seasonal patterns in phytoplankton photosynthetic parameters and primary production at a coastal NW Mediterranean site. Sci Mar 80S1: 63–77.

Gasol JM, Morán XAG. (2015). Flow cytometric determination of microbial abundances and its use to obtain indices of community structure and relative activity. Hydrocarb Lipid Microbiol Protoc - Springer Protoc Handbooks 1–29.

Gilbert JA, Steele JA, Caporaso JG, Steinbrück L, Reeder J, Temperton B, et al. (2012). Defining seasonal marine microbial community dynamics. ISME J 6: 298–308.

Gloor GB, Macklaim JM, Pawlowsky-Glahn V, Egozcue JJ. (2017). Microbiome datasets are compositional: And this is not optional. Front Microbiol 8: 1–6.

Grasshoff K, Ehrhardt M, Kremling K. (1983). Methods of seawater analysis. 2nd ed. Verlag Chemie, Weinheim.

Hatosy SM, Martiny JBH, Sachdeva R, STeele J, Fuhrman JA. (2013). Beta diversity of marine bacteria depends on temporal scale. Ecology 94: 1898–1904.

Hojerová E, Mašín M, Brunet C, Ferrera I, Gasol JM, Koblížek M. (2011). Distribution and growth of aerobic anoxygenic phototrophs in the Mediterranean Sea. Environ Microbiol 13: 2717–2725.

Kane MD. (2004). Microbial observatories: exploring and discovering microbial diversity in the 21st century. Microb Ecol 48: 447–448.

Kim DY, Countway PD, Jones AC, Schnetzer A, Yamashita W, Tung C, et al. (2014). Monthly to interannual variability of microbial eukaryote assemblages at four depths in the eastern North Pacific. ISME J 8: 515–530.

Koblížek M. (2015). Ecology of aerobic anoxygenic phototrophs in aquatic environments. FEMS Microbiol Rev 39: 854–870.

Koblížek M, Mlčoušková J, Kolber Z, Kopecký J. (2010). On the photosynthetic properties of marine bacterium COL2P belonging to *Roseobacter* clade. Arch Microbiol 192: 41–49.

Koblížek M, Moulisová V, Muroňová M, Oborník M. (2015). Horizontal transfers of two types of *puf* operons among phototrophic members of the *Roseobacter* clade. Folia Microbiol (Praha) 60: 37–43.

Koblížek M, Zeng Y, Horák A, Oborník M. (2013). Regressive evolution of photosynthesis in the *Roseobacter* clade. In: Vol. 66. Advances in Botanical Research. Elsevier, pp 385–405.

Kurtz ZD, Müller CL, Miraldi ER, Littman DR, Blaser MJ, Bonneau RA. (2015). Sparse and compositionally robust inference of microbial ecological networks. PLoS Comput Biol 11: 1–25.

Lami R, Cottrell MT, Ras J, Ulloa O, Obernosterer I, Claustre H, et al. (2007). High abundances of aerobic anoxygenic photosynthetic bacteria in the South Pacific Ocean. Appl Environ Microbiol 73: 4198–4205.

Legendre P, Legendre L. (1988). Numerical Ecology, Volume 24. Developments Environ Model 24: 870.

Lehours AC, Cottrell MT, Dahan O, Kirchman DL, Jeanthon C. (2010). Summer distribution and diversity of aerobic anoxygenic phototrophic bacteria in the Mediterranean Sea in relation to environmental variables. FEMS Microbiol Ecol 74: 397–409.

Lehours AC, Enault F, Boeuf D, Jeanthon C. (2018). Biogeographic patterns of aerobic anoxygenic phototrophic bacteria reveal an ecological consistency of phylogenetic clades in different oceanic biomes. Sci Rep 8: 4105.

Lehours AC, Jeanthon C. (2015). The hydrological context determines the beta-diversity of aerobic anoxygenic phototrophic bacteria in European Arctic seas but does not favor endemism. Front Microbiol 6: 1–9.

Logares R. (2017). Workflow for analysing MiSeq amplicons based on UPARSE v1.5.

Louca S, Parfrey LW, Doebeli M. (2016). Decoupling function and taxonomy in the global ocean microbiome. Science 353: 1272–1277.

Mahé F, Rognes T, Quince C, de Vargas C, Dunthorn M. (2014). Swarm: robust and fast clustering method for amplicon-based studies. PeerJ 2: e593.

Martin-Platero AM, Cleary B, Kauffman K, Preheim SP, McGillicuddy DJ, Alm EJ, et al. (2018). High resolution time series reveals cohesive but short-lived communities in coastal plankton. Nat Commun 9: 266.

Martin M. (2011). Cutadapt removes adapter sequences from high-throughput sequencing reads. EMBnet.journal 17: 10.

Martiny AC, Treseder K, Pusch G. (2013). Phylogenetic conservatism of functional traits in microorganisms. ISME J 7: 830–838.

Massana R, Murray AE, Preston CM, Delong EF, Massana R, Murray AE, et al. (1997). Vertical distribution and phylogenetic characterization of marine planktonic Archaea in the Santa Barbara Channel. Appl Environ Microbiol 63: 50–56.

McMurdie PJ, Holmes S. (2013). Phyloseq: An R package for reproducible interactive analysis and graphics of microbiome census data. PLoS One 8: e61217.

Oksanen J, Blanchet FG, Kindt R, Legendre P, Minchin PR, O’Hara RB, et al. (2013). Package ‘vegan’. Community Ecol Packag version 2. doi: 10.4135/9781412971874.n145.

Pedersen TL. (2017). ggraph: An implementation of grammar of graphics for graphs and networks. https://cran.r-project.org/package=ggraph.

Pedrós-Alió C. (2012). The rare bacterial biosphere. Ann Rev Mar Sci 4: 449–466.

R Core Team. (2014). R: A language and environment for statistical computing. https://www.r-project.org/.

Ritchie AE, Johnson ZI. (2012). Abundance and genetic diversity of aerobic anoxygenic phototrophic bacteria of coastal regions of the pacific ocean. Appl Environ Microbiol 78: 2858–2866.

Romera-Castillo C, Álvarez-Salgado XA, Galí M, Gasol JM, Marrasé C. (2013). Combined effect of light exposure and microbial activity on distinct dissolved organic matter pools. A seasonal field study in an oligotrophic coastal system (Blanes Bay, NW Mediterranean). Mar Chem 148: 44–51.

Salazar G, Cornejo-Castillo FM, Benítez-Barrios V, Fraile-Nuez E, Álvarez-Salgado XA, Duarte CM, et al. (2015). Global diversity and biogeography of deep-sea pelagic prokaryotes. ISME J 1–13.

Salter I, Galand PE, Fagervold SK, Lebaron P, Obernosterer I, Oliver MJ, et al. (2015). Seasonal dynamics of active SAR11 ecotypes in the oligotrophic Northwest Mediterranean Sea. ISME J 9: 347–360.

Shade A, Caporaso JG, Handelsman J, Knight R, Fierer N. (2013). A meta-analysis of changes in bacterial and archaeal communities with time. ISME J 754: 1493–1506.

Shade A, Jones SE, Caporaso JG, Handelsman J, Knight R, Fierer N, et al. (2014). Conditionally rare taxa disproportionately contribute to temporal changes in microbial diversity. MBio 5: 1–9.

Sieracki ME, Gilg IC, Thier EC, Poulton NJ, Goericke R. (2006). Distribution of planktonic aerobic anoxygenic photoheterotrophic bacteria in the northwest Atlantic. Limnol Oceanogr 51: 38–46.

Smith DC, Azam F. (1992). A simple, economical method for measuring bacterial protein synthesis rates in seawater using tritiated-leucine. Mar Microb Food Webs 6: 107–114.

Sogin ML, Morrison HG, Huber JA, Welch DM, Huse SM, Neal PR, et al. (2006). Microbial diversity in the deep sea and the underexplored ‘rare biosphere’. Proc Natl Acad Sci 103: 12115 LP-12120.

Sunagawa S, Coelho LP, Chaffron S, Kultima JR, Labadie K, Salazar G, et al. (2015). Ocean plankton. Structure and function of the global ocean microbiome. Science 348: 1261359.

Vila-Reixach G, Gasol JM, Cardelús C, Vidal M. (2012). Seasonal dynamics and net production of dissolved organic carbon in an oligotrophic coastal environment. Mar Ecol Prog Ser 456: 7–19.

Weiss S, Van Treuren W, Lozupone C, Faust K, Friedman J, Deng Y, et al. (2016). Correlation detection strategies in microbial data sets vary widely in sensitivity and precision. ISME J 10: 1–13.

Xia LC, Steele JA, Cram JA, Cardon ZG, Simmons SL, Vallino JJ, et al. (2011). Extended local similarity analysis (eLSA) of microbial community and other time series data with replicates. BMC Syst Biol 5: S15.

Yooseph S, Sutton G, Rusch DB, Halpern AL, Williamson SJ, Remington K, et al. (2007). The Sorcerer II global ocean sampling expedition: Expanding the universe of protein families. PLoS Biol 5: 0432–0466.

Yutin N, Suzuki MT, Béjà O. (2005). Novel primers reveal wider diversity among marine aerobic anoxygenic phototrophs. Appl Environ Microbiol 71: 8958–8962.

Yutin N, Suzuki MT, Teeling H, Weber M, Venter JC, Rusch DB, et al. (2007). Assessing diversity and biogeography of aerobic anoxygenic phototrophic bacteria in surface waters of the Atlantic and Pacific Oceans using the Global Ocean Sampling expedition metagenomes. Environ Microbiol 9: 1464–1475.

